# Deciphering PD1 activation mechanism from molecular docking and molecular dynamic simulations

**DOI:** 10.1101/2021.09.16.460652

**Authors:** Luis F. Ponce, Daniel P. Ramirez-Echemendia, Kalet Leon, Pedro A. Valiente

## Abstract

The activation of T cells is normally accompanied by inhibitory mechanisms within which the PD1 receptor stands out. Upon binding the ligands PDL1 and PDL2, PD1 drives T cells to an unresponsive state called exhaustion characterized by a markedly decreased capacity to exert effector functions. For this reason, PD1 has become one of the most important targets in cancer immunotherapy. Despite the numerous studies about PD1 signaling modulation, how the PD1 signaling is activated upon the ligands’ binding remains an open question. Several experimental facts suggest that the activation of the PD1-PLD1 pathway depends on the interaction with an unknown partner at the cellular membrane. In this work, we investigate the possibility that the target of PD1-PDL1 is the same PD1-PDL1 complex. We combined molecular docking to explore different binding modes with molecular dynamics and umbrella sampling simulations to assess the complexes’ stability. We found a high molecular weight complex that explains the activation of PD1 upon PDL1 binding. This complex has an affinity comparable to the PD1-PDL1 interaction and resembles the form of a linear lattice.

## Introduction

Persistent antigen stimulation leads the cells to an inactivated state called exhaustion (Yi, Cox, and Zajac 2010). In the transition to this state, T cells experience a progressive loss of effector functions and become unable to exert effective killing mechanisms allowing the establishment of chronic diseases such as viral infections and cancer (J. Lee et al. 2015). One of the major molecular fingerprints of the exhaustion state is the expression of inhibitory molecules among which programmed cell death (PD1) stands out (Blank and Mackensen 2007). Upon the binding of the ligands PDL1 and PDL2, PD1 inhibits mainly the cascade of the costimulatory molecules such as CD28 and the signaling through the T cell receptors (Hui et al. 2017).

The capacity of PD1 to inactivate T cells is used by tumors to escape from immunosurveillance (Alsaab et al. 2017). The PDL1 and PDL2 expressed in the tumor cells and surrounding tissues interact with the PD1 expressed in the activated T cells which promotes an immune tolerance where the immune system is incapable to mount a proper inflammatory response, even in the presence of activating antigens (Mahoney, Rennert, and Freeman 2015). Because of its inhibitory capacity, PD1 has become one of the major targets of cancer immunotherapies (Alsaab et al. 2017). Patients with lung cancers, melanoma, Hodgkin, and large B cell lymphomas have been the most beneficiated with anti PD1 drugs (Wu et al. 2019).

Despite the large number of studies on PD1 and PDL1, little is known about how the binding of PDL1/PDL2 induces the PD1 inhibitory signal. Some theoretical works suggest that the complex PD1-PDL1 has an unknown partner at the cell membrane (Ponce et al. in process). The results of Hui (Hui et al. 2017) suggest that CD28 costimulatory molecules and the T cell receptors as well as the same PD1-PDL1 complexes can be the membrane target of the PD1-PDL1 complex due to the co-clusterization in the immunological synapse region. The facts that the soluble form of PDL1 induces the PD1 signal (Liang et al. 2017; Frigola et al. 2011) and the soluble form of PD1 is capable to induce the PDL1/2 signaling in dendritic cells (Kuipers et al. 2006) suggest that the partner of the PD1-PDL1/2 complex in the cell membrane is the same complex.

In this work, we explore the possibility that the molecular partner of the PD1-PDL1 complex in the cell membrane is itself by using different techniques of computational structural biology. First, we used the docking protocol to find the most likely form of the interaction between two PD1-PDL1 complexes. The molecular dynamic (MD) and umbrella sampling simulations (US) were used to assess the stability of the different interaction forms. We found a tetrameric complex that is stable in the MD simulations and displays a comparable binding affinity with the PD1-PDL1 complex. A tetrameric complex for the murine PD1-PDL2 interaction with itself was built by structural alignment showing stability and comparable interaction energy, which suggests that this interaction is likely to occur in nature. This form of interaction corresponds to an oligomer of high molecular weight in the form of a linear lattice.

## Methods

### Structures preparation

The crystal structure of hPD1-hPDL1 and mPD1-mPDL2 complexes were obtained from the PDB database (PDB codes 4ZQK and 3BIK). Missing atoms were built by the method of Chinea (Chinea et al. 1995a) implemented in the *‘whatif’* server swift.cmbi.umcn.nl. Gaps in the structure of hPD1-hPDL1 were predicted with the method of Chys and Chacon (Chinea et al. 1995b; Chys and Chacón 2013) by using the server.rcd.chaconlab.org.

### Global and local docking with Rosetta

To carry out the docking protocol it was used the RosettaDock v3.2 (Gray et al. 2003) with RosettaScripts (Fleishman et al. 2011) considering only rigid body movements. Two docking classes were considered: DockingProtocol that generates 5×10^5^ structures and DockingLowRes that generate 10^3^ structures. All the parameters used for the protocol were set to the default ones.

### Molecular dynamic simulations

All MD simulations were carried out in gromacs version 2018 (Abraham et al. 2018) with periodic boundary conditions and the amber99sb force field (Lindorff-Larsen et al. 2010). All the simulations were carried out for 500ns. The protonation state of ionizable residues was estimated with the program PDB2PQR (Dolinsky et al. 2007), which uses PROPKA for PKA values prediction and is implemented in the server nbcr-222.ucsd.edu/pdb2pqr_2.1.1. The systems were solvated and neutralized (Na^+^/Cl^-^) by explicit water molecules, which were modeled by using the TIP3P parameters set (Jorgensen et al. 1983), in a rhombic dodecahedron box. The distance between the complexes and the edge of the box is 10 Å.

The systems were relaxed by 1000 steps using the steepest descent algorithm followed by 1000 steps by the conjugate gradient method, where the protein was held fixed by 10 kCal/mol Å constraint. After relaxation, systems were slowly heated to reach the temperature of 310 K. Newton’s equation of motion was solved using the Leapfrog scheme (Jorgensen et al. 1983; Verlet 1967), with an integration time step of 2fs. The temperature was controlled using a weak coupling to a bath with a time constant of 0.1 ps (Berendsen et al. 1984). For the pressure control, it was employed a Berendsen coupling algorithm with a time constant of 1.0 ps (Berendsen et al. 1984). In each replicate of the MD simulation, the Initial velocities were randomly generated following Maxwell distribution at 1 K, according to the atomic masses.

Motion equations were integrated with 2fs time step (Van Gunsteren and Berendsen 1988). The pressure was kept constant with the Parrinello-Rahaman coupling algorithm (Parrinello and Rahman 1981). A cutoff radii of 1.2 nm was established for the calculation of van der Waals (vdW) and short-range electrostatic interactions. The Particle Mesh Ewald (PME) summation (Essmann et al. 1995) was used to compute the long-range electrostatic interactions.

### Umbrella sampling

Umbrella sampling protocol (Torrie and Valleau 1977) was performed as indicated in the work of Lemkul and Bevan (Lemkul and Bevan 2010). The starting structures for the pulling protocol were taken directly from frames of the ordinary molecular dynamics. Pulling force was applied over α carbons of PD1, PD1-PDL1, and PD1-PDL2 complexes to separate them from PDL1, PD1-PDL1, and PD1-PDL2 complexes respectively.

The pulling force was applied toward z-direction without orthogonal restraints. Position restraints were considered over α-carbons of three residues of the fixed molecule or complex to avoid both rotations and translations of the systems. In the case of the simulation for the separation of PD1 from PDL1, these restraints were applied over 86I, 64I, and 125R of PDL1. In the case of the simulation separating PD1-PDL1 complexes we considered three position restraints over 57T, 91Q, and 124K in PDL1; and 51T, 79L, and 112L in PD1. In the case of the separation of mPD1-mPDL2 complexes, the position restraints were considered over T20, K45, and Q63 in PD1; and S39, Q60, and Y114 in PDL2. The force constant and the pulling rate were set to 1000 kJ /mol.nm^2^ and 10^−4^ nm/ps respectively. The frames for umbrella sampling windows were chosen to be separated in 1 Å.

The potential mean force (PMF) along the reaction coordinate was computed by the weighted histogram analysis method (WHAM) implemented in gromacs (Kumar et al. 1992; Hub, de Groot, and van der Spoel 2010). The last 100 ns of the MD simulation were considered for the calculation of the rotational and translational energies according to equation 11 of (M. S. Lee and Olson 2006).

## Results

### Docking

Rosetta global docking was carried out for 10^5^ structures. In these structures, one of the hPD1-hPDL1 complexes was fixed and the other was moved around to randomly generate the possible interaction modes (figure 1 panel A). The global docking procedure was carried out blindly without using any information about the previously predicted regions to be important for the interaction with the possible PD1-PDL1 partner (Ponce et al. 2021). The best 10 structures according to the score reported by the Rosetta program (figure 1 panel B) were selected for further analysis.

**Figure 1:**
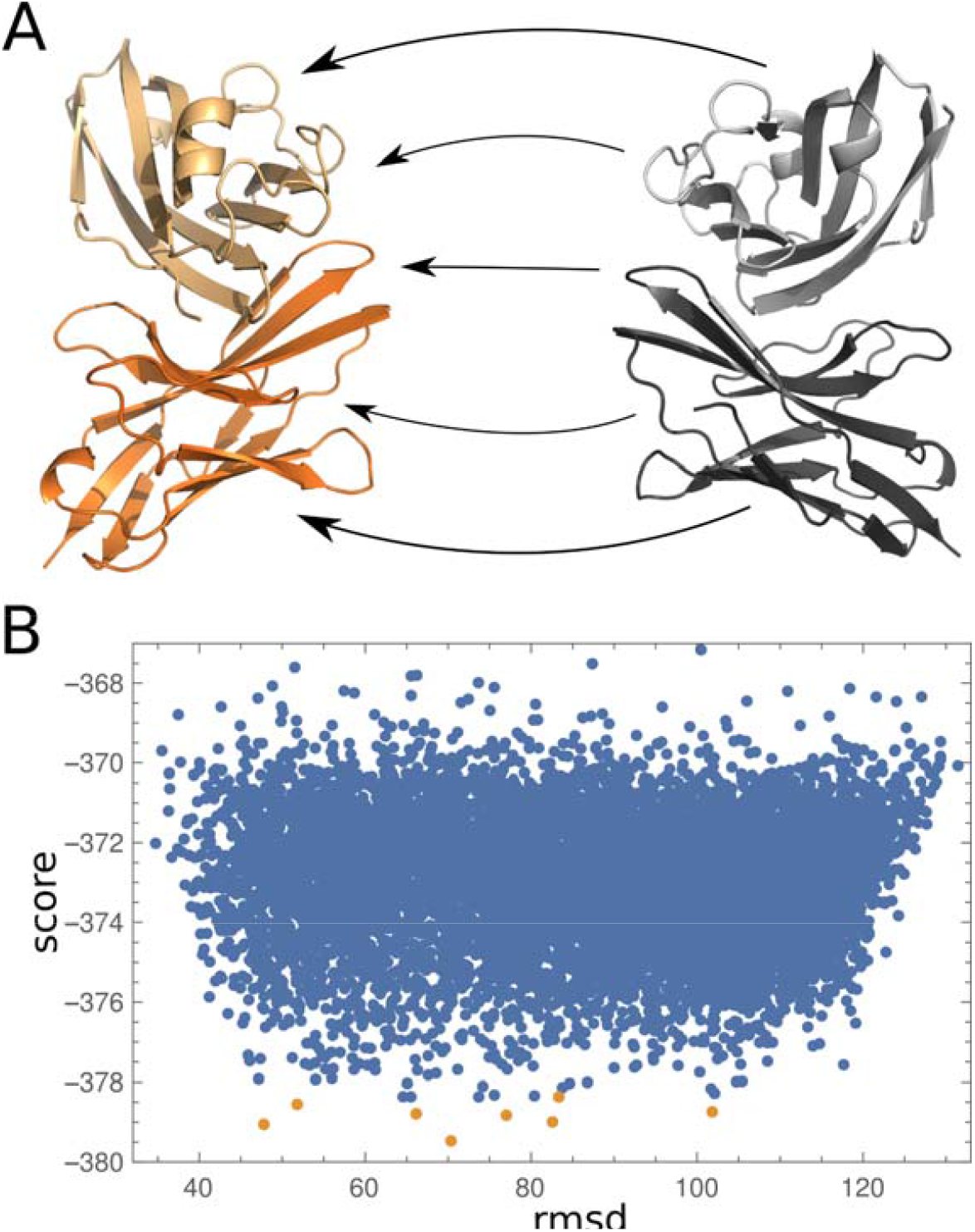
Global docking scheme for hPD1-hPDL1 complex interaction with itself. A) One of the PD1-PDL1 receptor complexes was fixed (the one in orange), while the other was moved around the first one exploring different interaction ways. The PDL1 structures are represented in lighter colors while darker colors correspond to the PD1 structures. B) Global docking scores for every analyzed structure for the interaction between PD1-PDL1 complex with itself. Each point corresponds to a different way of interaction. The best 10 structures determined by the score are represented in orange and were selected for further analysis.

Among the best 10 structures, those for which both PD1 and PDL1 belong to the interaction interface were selected for local docking assessment. Only 3 structures (called here tetramer T1-T3, figure 2 panels A-C) displayed a funneling pattern in the local docking suggesting that they can be significant biological structures (figure 2 panels D-F). The docking score was similar for these structures although slightly lower for the tetramer T1. The tetramers T2 and T3 correspond to asymmetric structures while tetramer T1 displays a clear translational symmetry where the interface of one of the complexes is completely exposed in the other one, allowing the binding of another/s complex/es.

**Figure 2:**
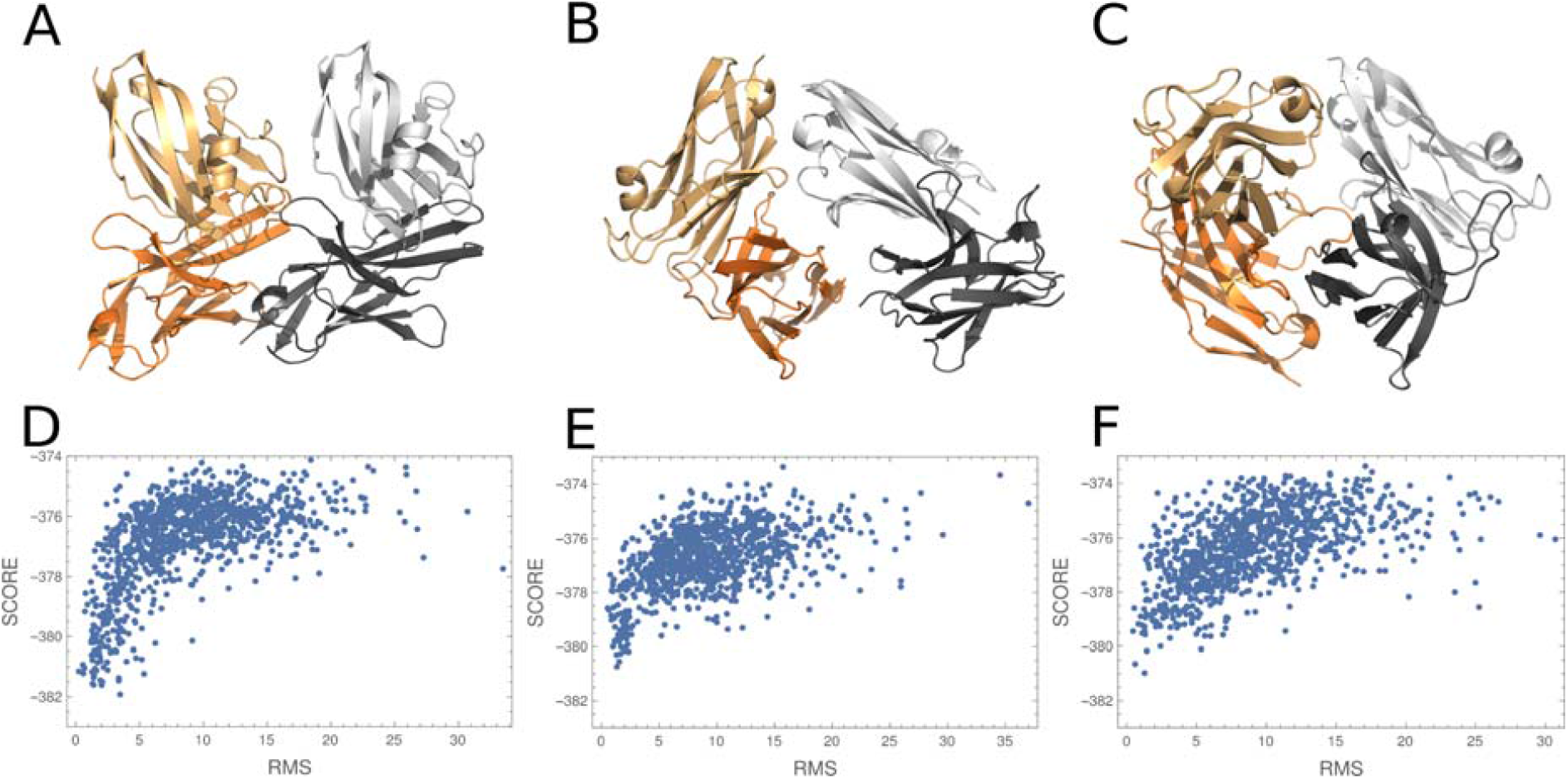
Local docking on the best interaction models between hPD1-hPDL1 complex with itself. A-C) Three structures (tetramers T1, T2, and T3 respectively) were selected among the best global docking results that displayed funneling patterns in the local docking. D-F Rosetta score for the structures fluctuating around the starting structures in local docking procedure. The RMSD value is measured in comparison with the structures coming from the global docking.

### MD simulations and free energy calculations

Although it seems that the first structure is the most appropriate one, it is important to note that docking results cannot be used for achieving conclusions especially for our case in which the interaction between the PD1-PDL1 complex with itself has not been experimentally demonstrated. For this reason, the MD simulations were used to assess the stability of the interactions in the predicted tetramers (figure 3).

**Figure 3.**
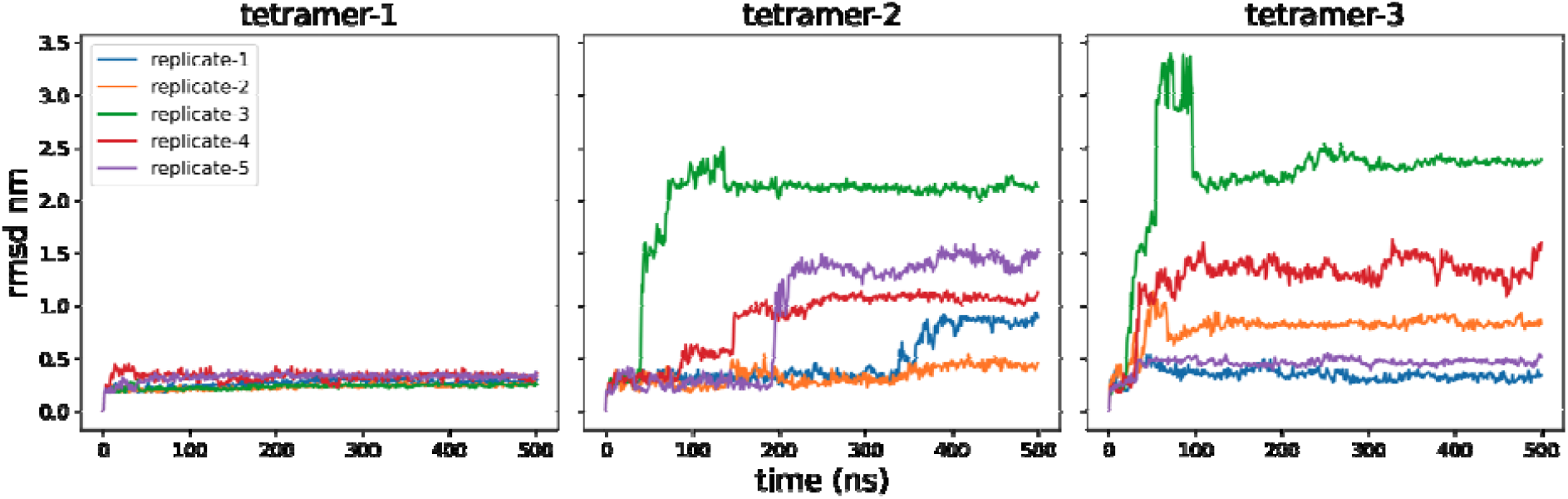
Instability of T2 and T3 in the molecular dynamic simulation. Graphs show the rmsd values on time for each MD replicate. Labels on the top indicate the tetramer model for each simulation

Five replicates of MD simulations were carried out for 500ns for a total of 2.5 µs. This time was enough to observe that the interactions described by tetramers T2 and T3 are unstable and lead the system to occupy other conformations (figure 4). On the other hand, in the simulations for the tetramer T1, the system remains stable during the time of the simulation in all the replicates which suggests that this tetramer is suitable to describe the PD1-PDL1 complex interaction with itself.

**Figure 3).**
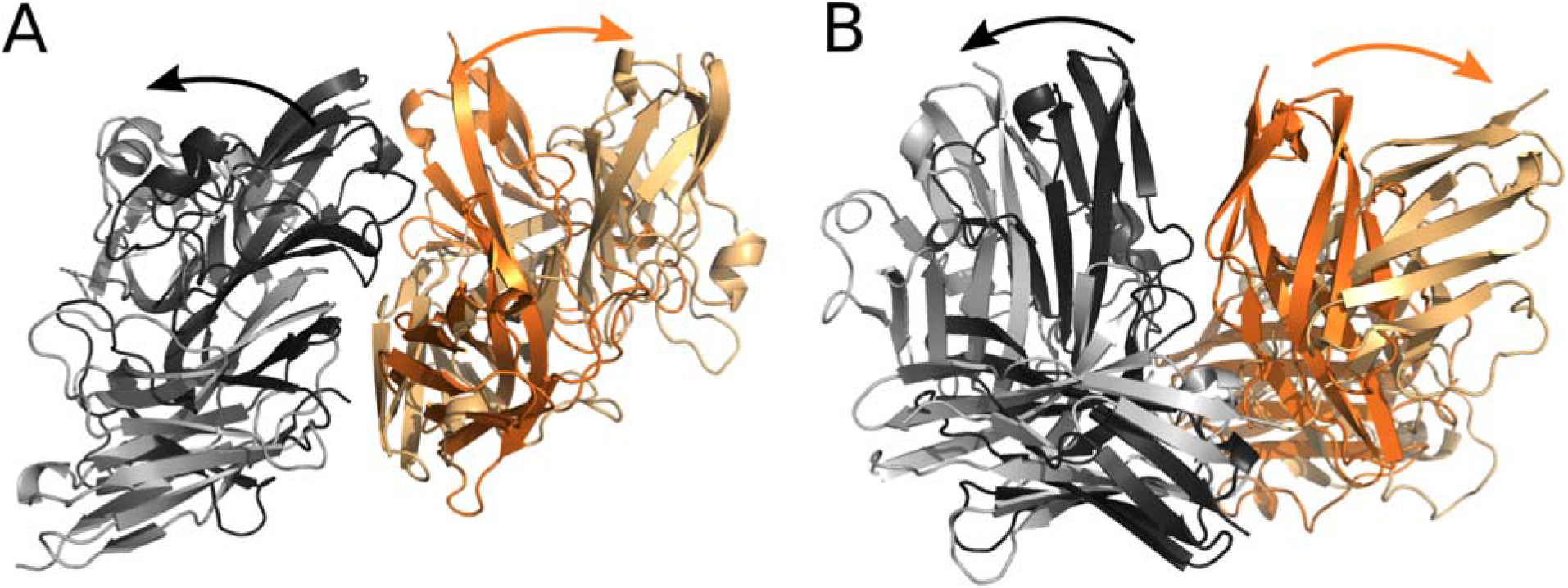
Instability in the kinetic of hPD1-hPDL1 interaction with itself in the T2 and T3 modes in the MD simulations. A, B) Example of separation of hPD1-hPDL1 from itself. Structures correspond to characteristic frames of the same MD simulation.

### Umbrella sampling

The stability observed in the replicates of MD simulation suggests that the complex may correspond to a biological structure. For this reason, we compare the strength of the interaction between PD1-PDL1 complexes with the one between PD1 and PDL1. To perform this comparison we carried out the umbrella sampling simulations for the separation of PDL1 from PD1 (the control) and the separation of one of the PD1-PDL1 complexes from the other one.

The separation of PDL1 from PD1 shows around -16.2 kcal/mol in the pmf value and around -9.2 kcal/mol after correcting the energy associated with translational and rotational degrees of freedom (figure 5 left panel and table 1). This result is consistent with the affinity estimated experimentally which is in the order of µM (Zhang et al. 2004). On the other hand, the separation of a PD1-PDL1 complex in the tetramer T1 shows a pmf value around -14.5 kcal/mol and around -10 kcal/mol after correcting the translational and rotational energies (figure 5 center panel and table 1). This interaction energy corresponds to an affinity of interactions of Kd below the µM range. This range for affinity values is more important for interactions occurring in the cell membrane where molecules are spatially restricted, which facilitates the interactions.

**Table 1:** Umbrella sampling results.

| Complex | PFM ± SD<br>(kcal/mol) | Rotational<br>(kcal/mol) | Translational<br>(kcal/mol) | Restraints<br>(kcal/mol) |
| --- | --- | --- | --- | --- |
| hPD1-hPDL1 | 16.17 (0.55) | -1.58 | -5.38 | -9.21 |
| *hPD1-hPDL1 | 14.52 (0.89) | -1.01 | -3.35 | -10.16 |
| *mPD1-mPDL2 | 15.01 (0.29) | -0.90 | -4.66 | -9.45 |

\* correspond to the separation from the complex in the tetramer T1

**Figure 5.**
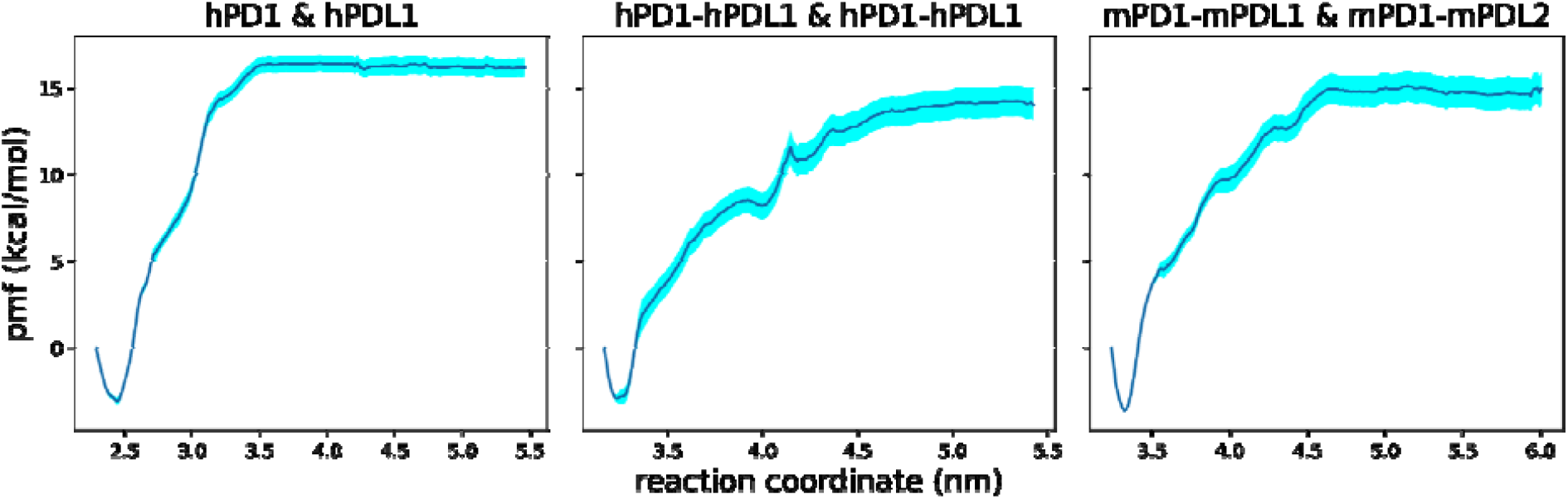
Umbrella sampling results for the separation of PD1-PDL1/2 T1 tetramers. Graphs show the behavior of the PMF value during the separation of the complexes along the reaction coordinate. Shadow areas around the mean value represent the standard deviation computed from 1000 bootstrap iterations. Labels at the top of each graph indicate the complex that is being separated.

To test how reliable was the interaction described by the tetramer T1, we used it as a mold for the interaction between the mPD1-mPDL2 complex with itself. The murine complex corresponds to two different molecules that are structurally similar to the human complex. The backrub protocol of rosetta (Lindorff-Larsen et al. 2010) was applied to the resulting complex after the alignment to solve the steric problems. A 500 ns MD simulation was carried out and a representative structure of the last 100 ns was used as the starting structure for the umbrella sampling simulation. The US simulation predicted a pmf value of -15 kcal/mole corresponding with -9.45 kcal/mole after correcting the translational and rotational degree of freedom associated energies (figure 5 right panel and table 1). This interaction energy is also consistent with the µM range of affinity.

### Main interactions description

The interaction mode between the PD1-PDL1 complexes in the tetramer T1 (figure 6 panel A) is consistent with the interaction mode of an antibody molecule which is consistent with our previous predictions (Ponce et al. in process). The major electrostatic interactions are established between the PD1 molecules (figure 6 panel B), which is consistent with the fact that PD1 signaling can be activated with different ligands, including the natural ligands PDL1 and PDL2 (Freeman et al. 2000; Latchman et al. 2001; Liang et al. 2017), which are similar in sequence and structure, as well as monoclonal antibodies that are completely different in both sequence and structure (Tyson 2010). The MD simulations showed three important interactions corresponding to hydrogen bonds or salt bridges that remain bound across most of the simulation time: the one formed between 61E and 139R (figure 6 panel C), between 101P and 143R (figure 6 panel D); and a salt bridge between 146E and 96R (figure 6 panel E).

**Figure 6.**
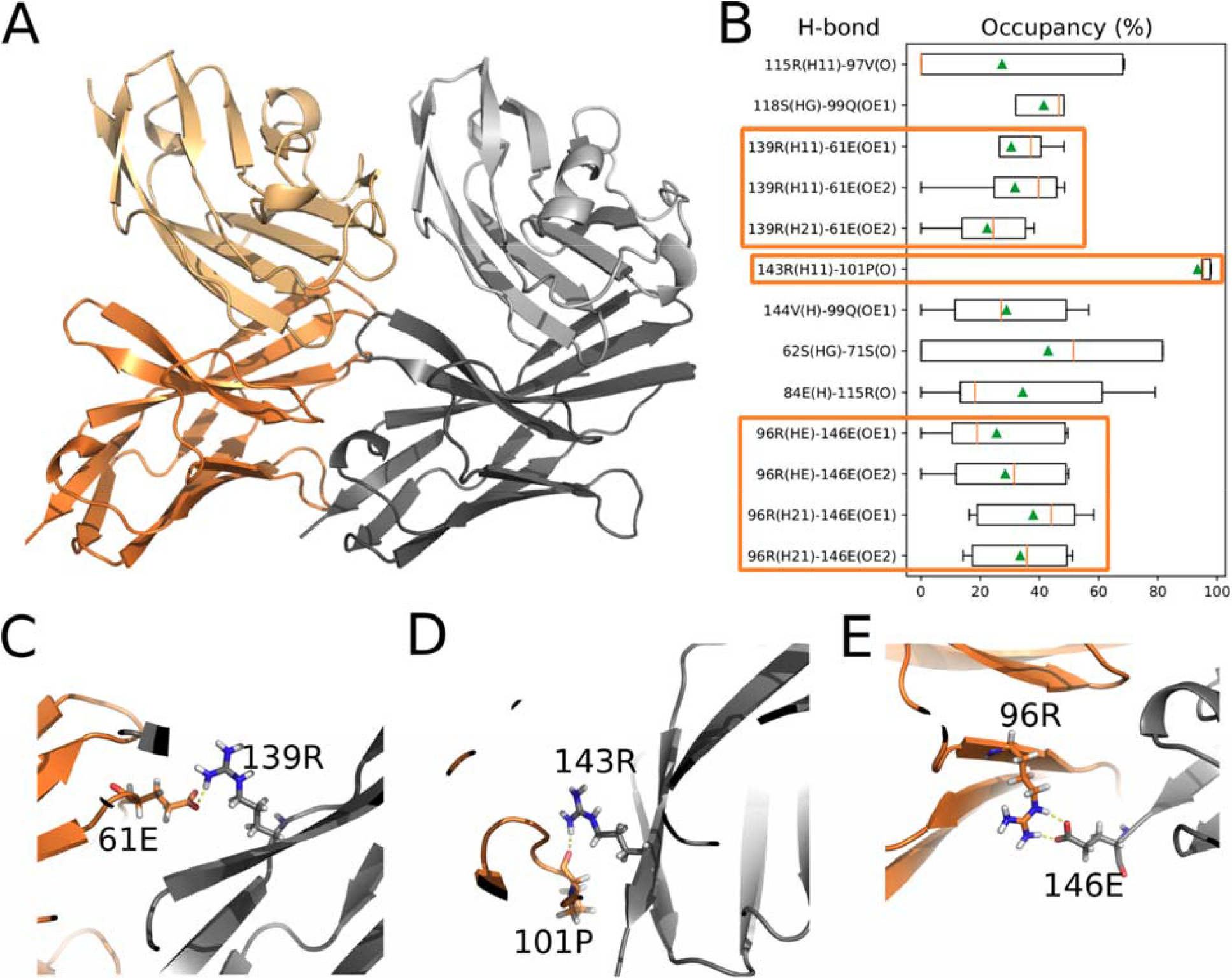
Main electrostatic interactions between PD1-PDL1 complexes in the tetramer T1. A) Structural representation of the tetramer T1. B) Occupancy of the main hydrogen bonds between PD1-PDL1 complexes across the simulation time considering the last 400ns of the five replicates of the MD simulations. Boxes indicate the statistics of occupancy (in percent) across the 5 replicates of MD simulations. Rectangles indicate the most stable interactions. Darker and lighter colors correspond to the PD1 and PDL1 molecules respectively. C-E) Representation of the main electrostatic interactions between PD1-PDL1 complexes in the T1 tetramer.

## Discussion

Despite the number of studies on PD1 interactions, the activation mechanism remains to be elucidated. It is well known that the binding of the ligands is translated in the recruitment of the activated SHP2 phosphatase intracellularly (Marasco et al. 2020). The main missing step in the PD1 activation mechanism is the connection between the binding of the ligands and the recruitment of the SHP2 enzyme. The fact that PD1 has a large and flexible tail between the topological and membrane domains makes it unlikely that the information about the binding of the ligands can be directly transmitted to the intracellular domain.

The results of Hui (Hui et al. 2017) showed that the main target for dephosphorylation of SHP2 enzyme is the CD28 costimulatory molecule, followed by the T cell receptor. However, the fact that CD80, the main ligand of CD28, binds to PDL1 in cis conformation and prevents the binding with PD1 (Zhao et al. 2019), makes unconvincing the idea of CD28 as the main PD1-PDL1/2 membrane target. On the other hand, several experimental results showed that in the immunological synapse, PD1 co-clusters with CD28 and TCR, which is consistent with the fact that these molecules are the major targets of PD1 inhibitory signal (Yokosuka et al. 2012; Hui et al. 2017). However, the activation of the cytosolic domain of PD1 and the recruitment of SHP2 does not require either CD28 or TCR intracellular domains (Hui et al. 2017).

The most clarifying work regarding the mechanism of the PD1 inhibitory signal is the one of Patsoukis (Patsoukis et al. 2020). In this work, it was found that the recruitment of SHP2 phosphatase activity depends on the interaction with at least two PD1 molecules and that the intracellular domains of PD1 interact upon the PDL1 binding. Joining the Patsoukis results with our predictions we may understand the full activation mechanism of PD1 inhibitory signaling (figure 7). Firstly, the ligands PDL1 and PDL2 bind to PD1 and induces the oligomerization of the extracellular domains. This oligomerization facilitates the interaction of the intracellular domains of PD1 and the further interaction with the activated SHP2 enzyme.

**Figure 7.**
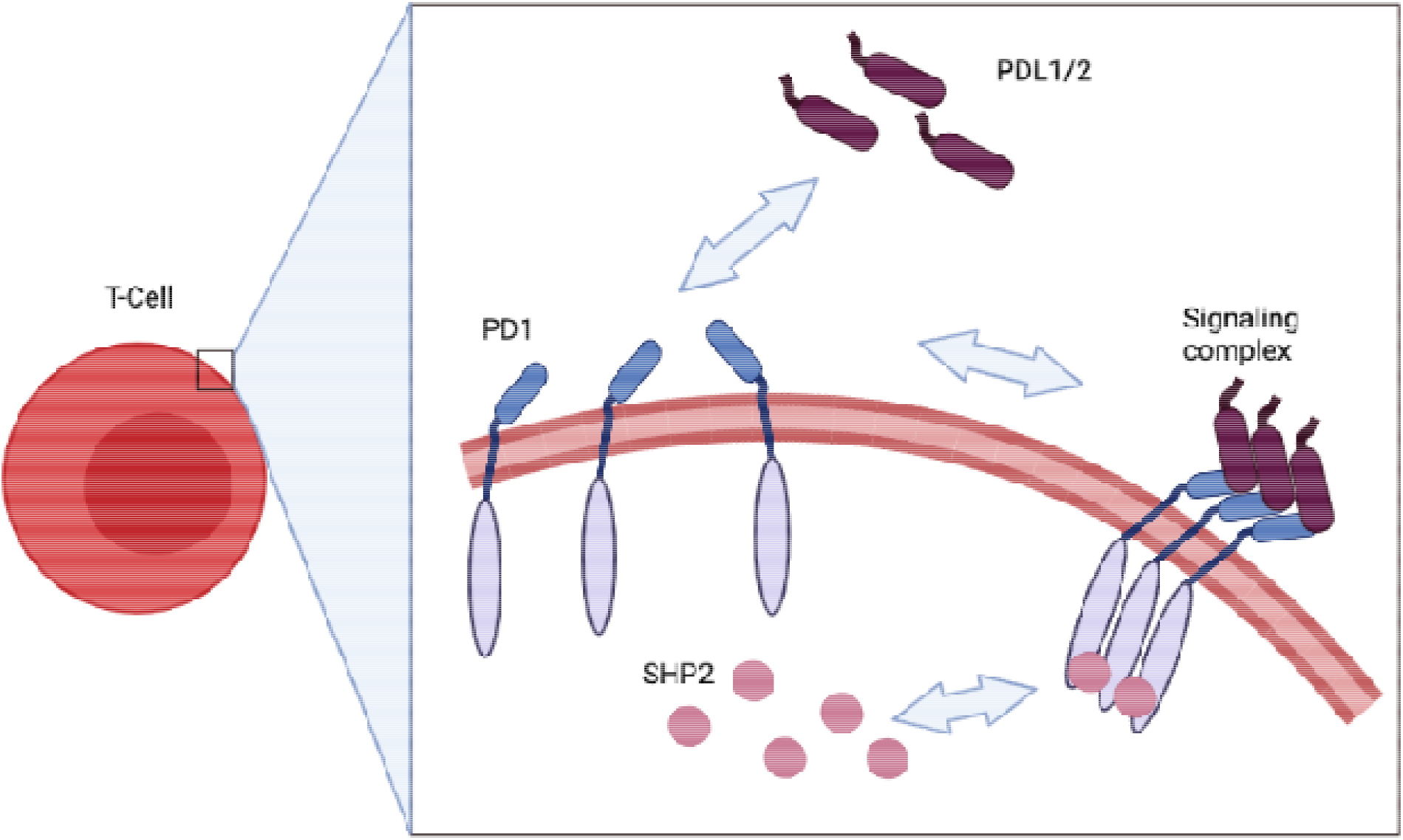
PD1 inhibitory signaling activation by PDL1 and PDL2

Our predictions are not only consistent with the results of Patsoukis (Patsoukis et al. 2020) but also with other experimental facts that indicate that the PD1 membrane target should interact more with the PD1 molecule. Furthermore, our predictions are consistent with our previous predictions for the relevance of the C’D and FG loops conformations for the PD1 signaling as well as the results of (Liu et al. 2017) for the CC’ loop conformational differences between bound and free PD1 states

One of the major limitations of the consequences deduced from our predictions is that large-size oligomers, are difficult to study experimentally, and may be confused with the precipitated fractions. Crystalizing the oligomers is also challenging, especially because of the filtration steps that may eliminate large-size oligomers. For this reason, crystal structures may correspond to those complexes that are not interacting. For example, the crystal structure of the human PD1-PDL1 complex (PDB 4zqk) contains PDL1 molecules with an Alanine residue in the first position that belongs to the signaling peptide. This Alanine induces a conformational change in the C’D region of PD1 that has an impact on the interaction with the other PD1-PDL1 complex according to our predictions.

Our results suggest new regions of PD1 that may be the target of anti-PD1 drugs, which should not prevent ligands’ binding. For example, the compounds explored in our previous work (Ponce et al. in process) can bind the PD1-PDL1 complex and prevent its binding to another PD1-PDL1 complex. The non-competitive PD1 inhibitors would be able to keep the PD1-PDL1/2 interaction and therefore, contribute to the cell-to-cell interaction, an interesting advantage over the competitive inhibitors that can improve the anticancer immunotherapies.

